# Speciation driven by hybridization and chromosomal plasticity in a wild yeast

**DOI:** 10.1101/027383

**Authors:** Jean-Baptiste Leducq, Lou Nielly-Thibault, Guillaume Charron, Chris Eberlein, Jukka-Pekka Verta, Pedram Samani, Kayla Sylvester, Chris Todd Hittinger, Graham Bell, Christian R Landry

## Abstract

Hybridization is recognized as a powerful mechanism of speciation and a driving force in generating biodiversity. However, only few multicellular species, limited to a handful of plants and animals, have been shown to fulfill all the criteria of homoploid hybrid speciation. This lack of evidence could lead to the misconception that speciation by hybridization has a limited role in eukaryotes, particularly in single-celled organisms. Laboratory experiments have revealed that fungi such as budding yeasts can rapidly develop reproductive isolation and novel phenotypes through hybridization, showing that in principle homoploid speciation could occur in nature. Here we report a case of homoploid hybrid speciation in natural populations of the budding yeast *Saccharomyces paradoxus* inhabiting the North American forests. We show that the rapid evolution of chromosome architecture and an ecological context that led to secondary contact between nascent species drove the formation of an incipient hybrid species with a potentially unique ecological niche.

**One Sentence Summary:** Chromosomal rearrangements and hybridization between two yeast lineages drive hybrid speciation after secondary contact.

## Introduction

Hybridization is a major force in evolution because it can prevent population divergence by maintaining gene exchange ^1^. However, hybridization can also lead to speciation by combining genomes that have been evolving independently and that, when combined, provide advantageous phenotypes to the hybrid individuals and populations ^2,3^. The genomic composition of the hybrid individuals may also lead to reproductive isolation with their parental lineages ^4^. The identification of homoploid hybrid speciation events, which occur without change in chromosome number – contrary to what is observed in allopolyploid and homopolyploid speciation – requires that we show the reproductive isolation of the hybrid with the parental species, the identification of traces of past hybridization in the genome, and that the isolating mechanisms are at least partly derived from hybridization ^5^. Despite extensive investigations in sexually reproducing microbes, no case of such homoploid speciation has been reported in these eukaryotes ^5^. Yet, laboratory experiments have shown that hybridization can contribute to rapid species formation in yeast ^6^.

Here we examine the genomics and the ecological bases of species formation in a yeast natural system. The budding yeast *Saccharomyces paradoxus* is the closest known relative of the model yeast *Saccharomyces cerevisiae. S. paradoxus* is a free-living saprophyte mostly found in the sap and on the bark of deciduous trees and their associate soil ^7-10^. *S. paradoxus* has a nearly worldwide distribution but contrary to *S. cerevisiae,* there is no evidence for its domestication by humans ^8,11,12^. Accordingly, the biogeography ^12^ and the genome evolution ^13^ of this unicellular fungus are mostly influenced by natural processes. Four genetically and phenotypically distinct lineages of *S. paradoxus* have been identified so far and correspond to populations from Europe, Far East Asia, America and North-East America ^14-16^. In North America, three lineages of *S. paradoxus* occur in partial sympatry. One of these lineages, corresponding to the European population (Lineage *SpA*), has a sparse distribution and strains of this lineage are mostly isogenic, due to a recent colonization event ^15^. The two other lineages in North America (*SpB* and *SpC*) are indigenous and are incipient species ^17^ differentially distributed along a southwest to northeast gradient (Figs. 1a, S1). *SpB* strains display enhanced fitness at high temperature and survival to a freeze-thaw cycle when compared to *SpC* strains, consistent with adaptations to climatic conditions ^16^. Despite the overlapping distributions and partial post-zygotic reproductive isolation between *SpB* and *SpC* ^17^, no first-generation hybrid has been identified so far, suggesting, along with with the monophyly of these lineages, that speciation has been initiated.

**Figure 1.**
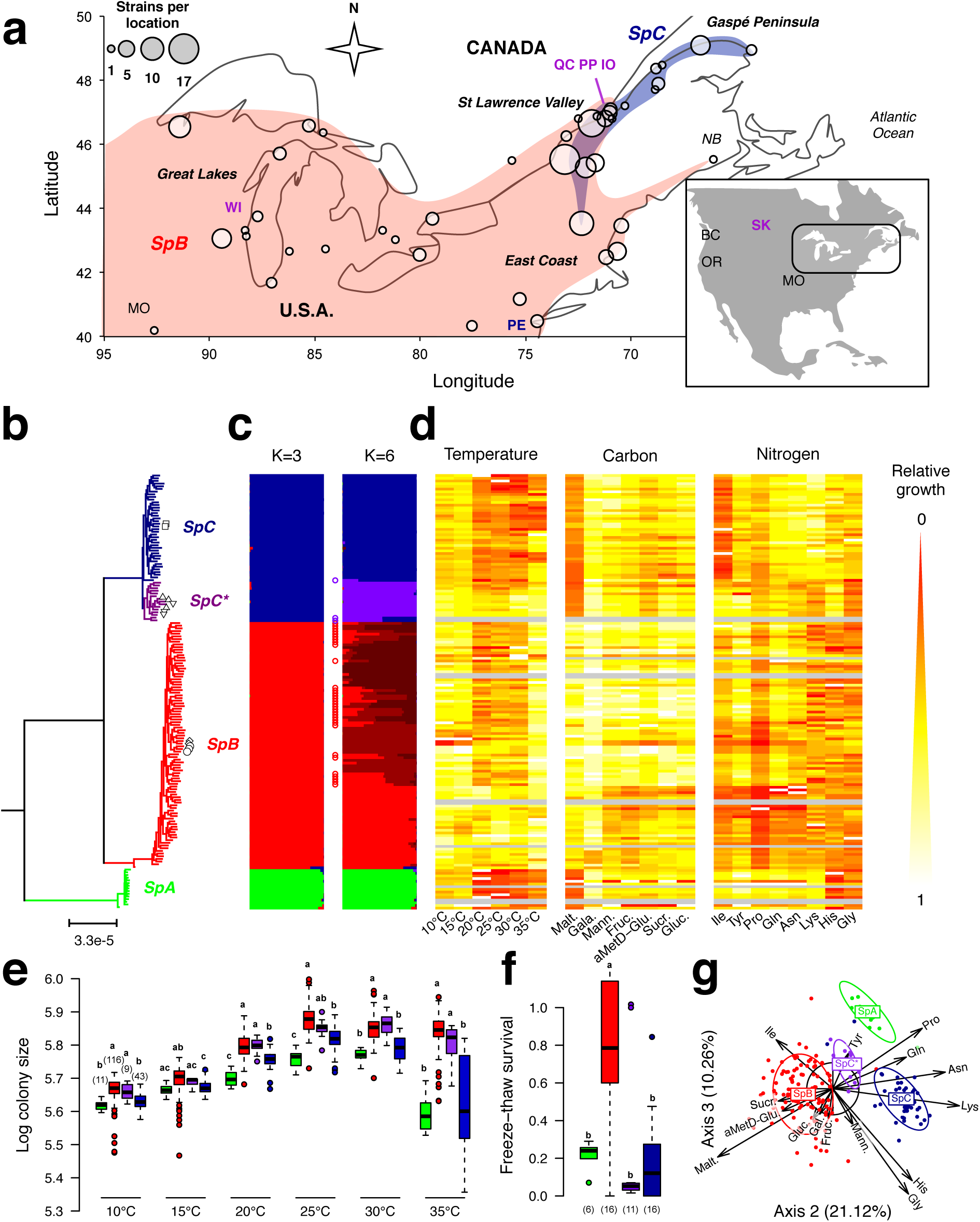
A cryptic *S. paradoxus* lineage revealed by a population structure and a distinct ecological niche from its sympatric close lineages. **a,** Sampling locations (circles) and distribution (inner frame) of *SpB* (red), *SpC* (dark blue; one strain in Pennsylvania) and the cryptic lineage *SpC*\* (purple). Location abbreviations are listed in Table S1. **b**, Eleven *SpC*\* strains form a monophyletic group within the *SpC* lineage. Phylogenetic tree of 161 strains rooted with *S. cerevisiae* (14,974 SNPs). *SpA* is the European lineage (green). Open symbols indicate replicates and spores from a same strain. **c**, STRUCTURE diagrams (6,881 randomly sampled SNPs, assuming K=3 and 6 populations) reveal 33 admixed *SpB* strains between three populations (<80% of assignation; red circles) and only three between *SpC* and *SpC*\* (PE and SK strains; purple circles). **d,** Relative growth (log of colony size; warm colors) measured at different temperatures and in limiting nutrient conditions for seven carbon and eight nitrogen sources for 151 strains. Ten strains (grey) did not pass the selection step. **e,** *SpC\** shows enhanced growth at high temperature, similarly to *SpB*. Bold letters indicate significant differences among lineages (Tukey test: *p*≤0.05). The number of strains per lineage is indicated. Error bars indicate interquartile ranges. **f,** Fraction of cells surviving a freeze-thaw cycle. *SpC\** and *SpC* show reduced survival as compared to *SpB* (*p*≤0.05). **g,** Metabolic profiles in limiting nutrient conditions summarized by a principal component analysis (axes 2 and 3 displayed). *SpC\** is intermediate between *SpC* (amino-acid preference) and *SpB* (carbon preference).

## Incipient speciation in wild yeast

We sequenced the genomes of 161 strains from the north east of North America to uncover the evolutionary history of this ongoing speciation event, using the European lineage (*SpA)* recently introduced in America as outgroup (see sup information section 1; Table S1). A phylogeny based on 14,974 filtered polymorphic sites confirms that *SpB* and *SpC* form distinct lineages with 2.09±0.01 % nucleotide divergence (Fig. 1b) and we observed no first generation hybrid among lineages, as supported by overall low heterozygosity (~0.1%; Fig. S2). The monophyletic *SpB* clade shows population substructure (Figs. 1b-c), consistent with its broad geographic distribution (Figs. 1a, S1). This analysis also revealed a third clade that is sister to the *SpC* lineage, which we call *SpC\** (Fig. 1b). *SpC\** consists of eleven *SpC* strains (22% of all sequenced *SpC*) showing limited genetic admixture with other *SpC* strains (Fig. 1c) and a narrow geographic distribution (Figs. 1a, S1).

Because *SpC*\* strains are mostly found near the southwest limit of the *SpC* distribution, this new lineage may be locally adapted and thus show increased fitness over *SpC* in the relatively warmer temperatures encountered in this region ^18^. We measured the growth of 182 strains along a temperature gradient from 10 to 35°C (Fig. 1d) and found that *SpC*\* strains show a growth advantage over *SpC* (+3-42% colony growth) similar to the *SpB* advantage over *SpC*, particularly at high temperature (+5-36%; *p*< 0.001; Tukey test; Tables S2-S3; Fig. 1e). We previously reported that populations of *S. paradoxus* showed variation for resistance to freeze-thaw cycles that is correlated with their latitude, with higher resistance in the south where these cycles are more frequent ^16^. We found that like *SpC*, *SpC*\* is highly sensitive to a freeze-thaw cycle (8.9% and 5.2% survival, respectively), while *SpB* has significantly higher survival (74.4%; *p* < 0.05; Tukey test; Tables S2-S3; Fig. 1f). We also examined other growth conditions that may reflect the large diversity of substrates on which yeasts were isolated^7,16,19^. Because *SpC* and *SpB* were recently shown to perform differently in maltose, lyxose and glucosamine ^21^, we first examined growth in various carbon sources that are known to shape sap microbial diversity ^20^. We also examined growth performance on different nitrogen sources, including several amino acids, which are known to be a major nitrogen source in tree sap ^22^. We found that the three lineages have contrasted metabolic profiles, as suggested by the growth advantage of *SpB* and *SpC* on specific carbon and nitrogen sources, respectively (Tukey test: *p*<0.001; Figs. 1g, Tables S2-S3). Overall, *SpC*\* strains show a metabolic performance profile intermediate between *SpB* and *SpC*. Assuming that yeasts have a have high dispersal potential like many other eukaryotic microbes^23^, these results suggest that the overall performance of *SpC\** in specific climatic conditions and substrates is a potential cause for its limited ecological distribution in the middle of the *SpB-SpC* range.

The monophyly of *SpC*\* and its contrasted phenotypes imply limited gene flow with *SpC*, despite these two lineages have overlapping distributions and co-occur on neighboring trees in the region of sympatry (Fig. S1). This indicates that *SpC*\* is reproductively isolated from *SpC* and could therefore represent an incipient species. Budding yeasts of the genus *Saccharomyces* have no obvious pre-zygotic isolation mechanisms in the laboratory^24^, which allows to measure progeny viability between *SpC*\*, *SpB* and *SpC* (Fig. 2). Our results indicate that survival of spores from *SpC*×*SpC*\* crosses is significantly lower than for *SpC*×*SpC* or for *SpC\**×*SpC*\* (*p*<0.001; Tukey test) and on the same order of magnitude as what is observed for *SpB*×*SpC*, and *SpB*×*SpC*\* (Fig. 2, Tables S4-S5), despite the nearly 10-fold difference in terms of molecular divergence (0.27% *vs*. 2.09%).

**Figure 2.**
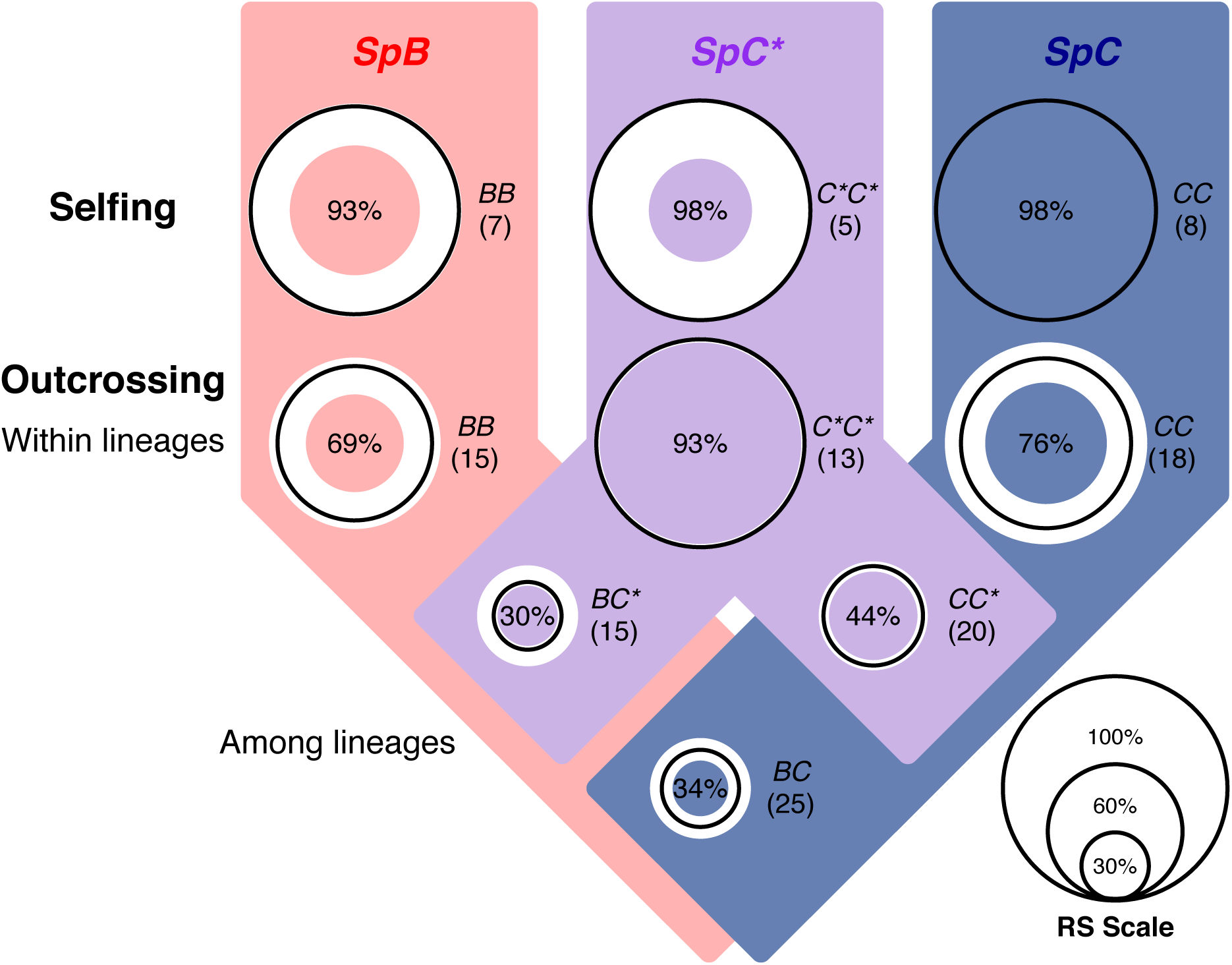
The *SpC\** lineage is an incipient species, as revealed by its limited reproductive success with its sister lineages. Post-zygotic reproductive success (RS) within and between lineages as estimated by progeny viability. Circle diameter is proportional to median RS (scale on bottom right; 75 percentiles of RS in white). The number of crosses per category is indicated. *SpC*\* shows partial reproductive isolation with *SpC* and *SpB,* to an extent that is similar to crosses between *SpC* and *SpB.*

## Genome dynamics following hybridization

We found that the *SpC\** lineage genome is a mosaic of *SpC* and *SpB* genotypes, with genomic islands of *SpB*-like alleles (i.e. fixed within *SpB*) present in *SpC*\* strains but absent in lineage *SpC* (Fig. 3a). These *SpB*-like regions correspond to 12 large regions (28-123kb) of high diversity within the *SpC*+*SpC\** clade (*He* > 0.005; *p* <0.0001; Figs. S3a-c), overlapping with 23 large regions (22-119kb) for which divergence between *SpB* and *SpC*+*SpC\** populations is low (*F*_ST_ < 0.75; *p* <0.0001; Figs. S3d-g) relative to the genome-wide average (*Fst* ≈ 1). These *SpB-*like regions most likely result from the introgression of *SpB* into *SpC*, as shown by a window-based phylogenetic analysis^25^ of 24 strains with high-quality genomes (Figs. S3h, S4a) and by a site-wise clustering analysis^18^ of the 161 strains (HC and LC combined; Fig. 3a). The monophyly of the *SpC*\* clade is maintained even when these introgressed regions are removed (Fig. S4b), suggesting that divergence, although very limited, has accumulated between *SpC* and *SpC*\* outside of the introgressions. The *SpB*-like regions represent 2.2-5.8% of the *SpC*\* genome, with 1.6% shared among all *SpC*\* strains. The common set of introgressed regions could be the remnants of a single ancestral hybridization event between the *SpB* and *SpC* lineages (marked as H0; Fig. 3a) and the polymorphic introgressed regions could result from the ongoing loss of *SpB*-like regions in *SpC*\* or from secondary introgression events (Table S6).

**Figure 3.**
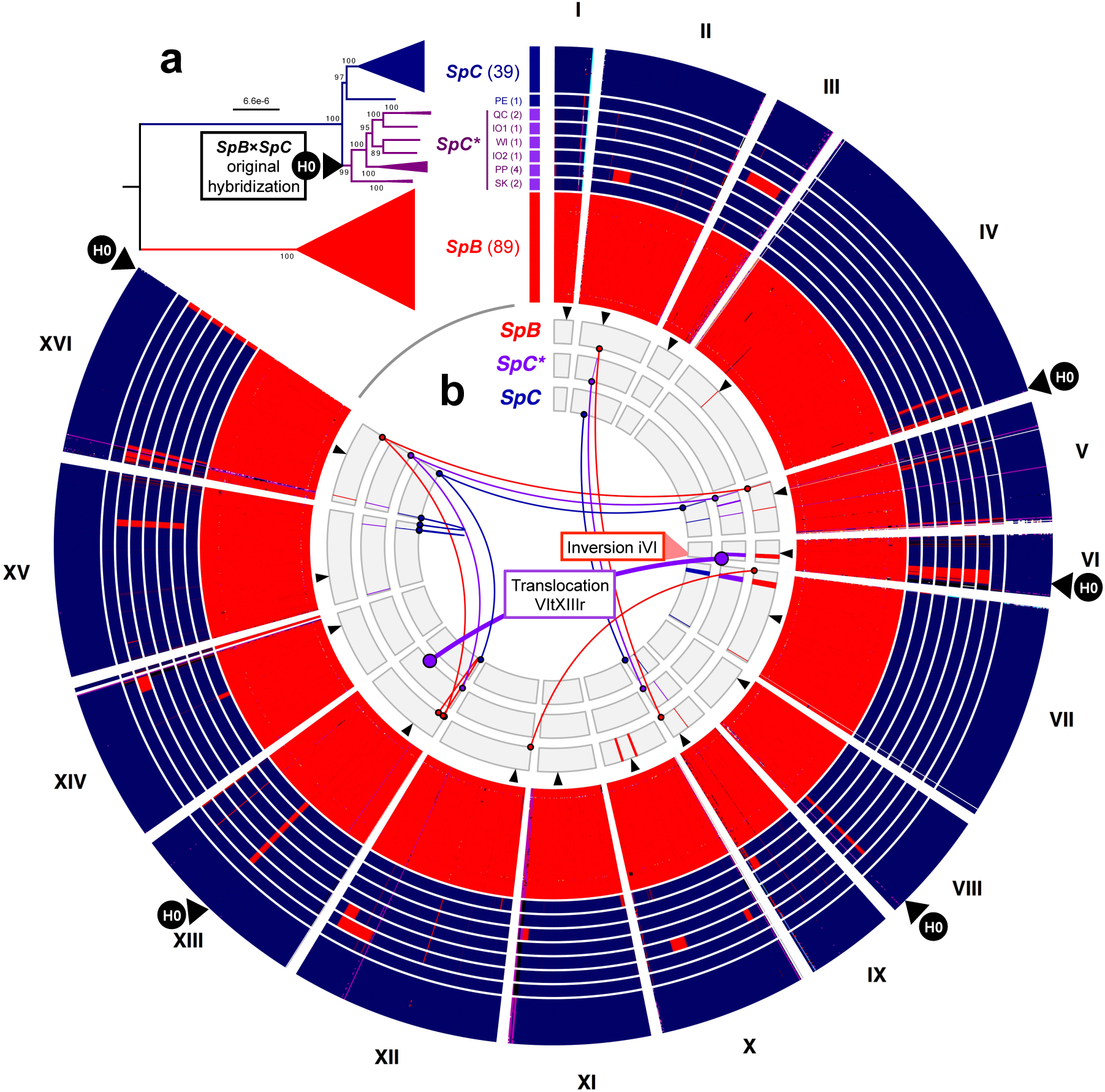
The *SpC*\* lineage is a mosaic of *SpC* and *SpB* genomes and results from past hybridization. **a,** *SpB*-like (red) and *SpC*-like (blue) genotypes (site-wise clustering approach, the *SpA* outgroup was removed) in 5kb windows along the genome of 147 *S. paradoxus* strains reveal that *SpC*\* strains are a mosaic of the two lineages, with a vast majority of *SpC* genotypes. *SpC*\* strains share a common set of *SpB*-like regions likely acquired during an original hybridization event between *SpB* and *SpC* (H0). Polymorphic *SpB*-like regions distinguish six *SpC*\* groups (see Table S6) that result from the ongoing loss of *SpB*-like regions or from secondary introgressions. **b,** Lineage-specific karyotypes of 19 strains with high-quality genomes (*SpA* removed). Telomere exchanges (thin lines), translocations (thick lines) and inversions (colored portions in chromosomes) as compared to *S. cerevisiae*, mapped on circular diagrams of the 16 chromosomes (I-XVI). A translocation (VItXIII) between chromosome VI and the right arm of chromosome XIII (240kb) is fixed within *SpC*\* and absent elsewhere. The 42kb inversion in chromosome VI overlapping with a *SpB*-like region (iVI) was likely transmitted from *SpB* to *SpC*\* and is absent among the *SpC* high-quality genomes (see also Fig. S7).

We examined chromosome size variation within and between the *SpC* and *SpC\** populations (Fig. S5) and found that the *SpC*\* karyotype was significantly different from both *SpC* and *SpB* based on chromosome migration profiles (*p* < 0.001; Tukey test; Fig. S6). Using *de novo* genome assembly, we found extensive variation in chromosome configuration among lineages (Fig. 3b) that supports the chromosome profile analysis (Figs. S5-S6). The patterns of inversion and smallscale exchange (≤ 20kb) among telomeric regions that are fixed within lineages confirm the overall *S. paradoxus* phylogeny (Fig. S7a). One exception is a 42kb inversion within chromosome VI (iVI; Fig. 3b). This inversion is only shared between *SpB* and *SpC*\* and is absent in *SpC* (Figs. S7a, Table S1). Another exception is the fusion between chromosomes VI and the right arm of chromosome XIII (VItXIII) that is absent in *SpC* but fixed in *SpC*\* (Figs. 3b, S7b). This fusion segregates at low frequency in *SpB* (3.3%; referred as *SpBf* strains; Figs. S7c). A phylogeny based on the iVI region reveals that *SpBf* is the sister group of *SpC*\*, suggesting that *SpC*\* inherited iVI and VItXIII from *SpBf* strains (Figs. S7d). Chromosomal rearrangements are often under-dominant and are thus unlikely to fix within a population^26^, which could explain the low frequency of VItXIII in *SpB*.

## Chromosomal changes and reproductive isolation

Given the low level of divergence between *SpC* and *SpC\** outside of the introgressed regions, reproductive isolation is unlikely driven by molecular incompatibilities such as mismatch repair^6^, although those could be important in *SpB*×*SpC* and *SpB*×*SpC*\* hybrids. We thus hypothesized that chromosomal changes originating from hybridization contribute to the reproductive isolation between *SpC*\* and *SpC*. Accordingly, these changes would miss-segregate in the progeny of the F1 hybrids and correlate with spore viability^27^ (Fig. 4a). We tested this hypothesis by sequencing 325 haploid F2 strains that we obtained from the sporulation of eight *SpC* × *SpC*\* diploid F1 hybrids (Fig. 4b, Table S7). We identified large genomic regions (100-200kb) where *SpC\** genotypes segregate unevenly in meiotic products with low spore viability (25%; Fig. 4c), corresponding to either *SpB*-like regions (IIL, VIr and XIIr; Fig. 4d) or to regions in linkage with the translocation (LT regions; Fig. 4e). We also observed systematic ploidy increase in the translocated arm of chromosome XIII (XIIIr; Fig. 4e). We confirmed that the genomic regions involved in at least two major chromosomal rearrangements that differentiate *SpC*\* from *SpC*, namely the inversion in chromosome VI (iVI) and the right arm of chromosome XIII (XIIIr), segregate unevenly in the surviving spores and contribute to reduced spore viability, as predicted from their simple Mendelian segregation (Fig. 5a). The iVI inversion also co-localizes with a *SpB*-like region that is preferentially transmitted in surviving spores (Fig. S8a). Both the frequency of the iVI inversion typical of *SpC*\* (Fig. 5b) and the ploidy of the XIIIr region (Fig. 5c) significantly decrease with spore survival (1,000 random permutations; *p*<0.05), showing that surviving spores systematically inherit these regions more than expected under normal segregation. The breakpoint of ploidy change in XIIIr coincides with the fusion point with chromosome VI in *SpC*\* (Fig. S8a), indicating that VItXIIIr causes lethal aneuploidy in *SpC* × *SpC*\* haploid hybrids by the sorting of XIIIr^27^ (Fig. 5a). We observed no significant ploidy variation on chromosome VI, suggesting that the over-representation of the iVI inversion in the spores and of the associated *SpB*-like region results from their linkage with XIIIr (LT region; Fig. S8b). These results show that the chromosomal polymorphism segregating at low frequency within *SpB* (VItXIIIr) was transmitted by hybridization with *SpC* and now causes reproductive barriers between *SpC\** and *SpC.* This transmission led to or co-occurred with the formation of the new lineage *SpC\** and contributed to reproductive isolation between *SpC\** and its parental lineage *SpC*. Because we observed similar patterns of abnormal segregation in other genomic regions (Fig. 5c), we reasoned that additional chromosomal differences between *SpC* and *SpC*\* also contribute to reproductive isolation (Figs. S6; see also supplementary material, section 27 for discussion). We estimated that taken together, chromosomal differences may lead to lethal aneuploidy in 45% of spores, hence contributing to up to 80% of actual reproductive isolation between *SpC* and *SpC*\* (Fig. 5a).

**Figure 4.**
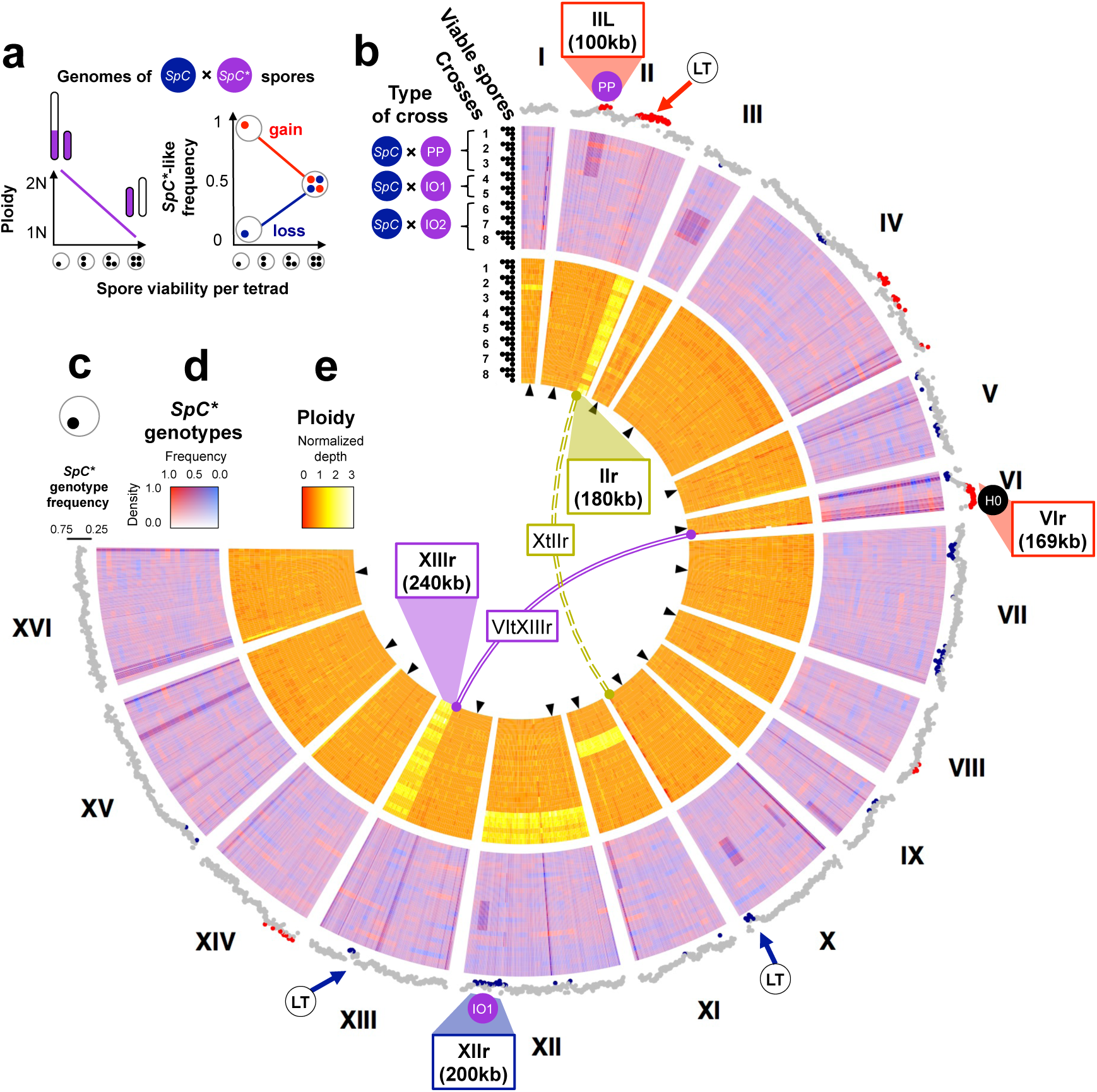
Chromosomal rearrangements and introgressed regions unevenly segregate in the *SpC*×SpC* hybrid progeny. **a,** Expected patterns of ploidy increase (left) and *SpB*-like genotype frequency variation (right) in genomic regions involved in decrease of *SpC*×SpC* spore viability. **b,** Genome sequencing of 384 spores from eight *SpC*×SpC* crosses (*SpC*\* types indicated) pooled according to the number of surviving spores per tetrad (*n*=1-4). **c,** Mean *SpB*like genotype frequency in tetrads with low viability (*n*=1) in discrete 20kb windows along the genome. Strong segregation biases toward *SpC* (blue) or *SpC\** (red) genotypes are observed in some *SpB*-like regions (IIL, VIr, XIIr) and regions linked to chromosomal translocations (LT). **d,** Frequency of *SpC*\* genotypes (red-blue heatmap, 5kb windows) per pool of spores. Dark regions (intensity proportional to divergence between *SpC* and *SpC*\* parents) correspond to *SpB*-like regions. **e.** Heatmap of ploidy variation (warm color scale, 5kb windows) suggests aneuploidy in hybrid progeny caused by chromosomal translocations. The rightmost breakpoint in XIIIr corresponds to the fusion point with chromosome VI (VItXIIIr, purple). A similar pattern in chromosome II (IIr) suggests the contribution of another chromosomal change (XtIIr, yellow; see Figs. S6, S8). The iVI inversion has no visible effect on ploidy. Ploidy was calculated as sequencing coverage per pool normalized by coverage in a control tetrad (*n*=4; removed).

**Figure 5.**
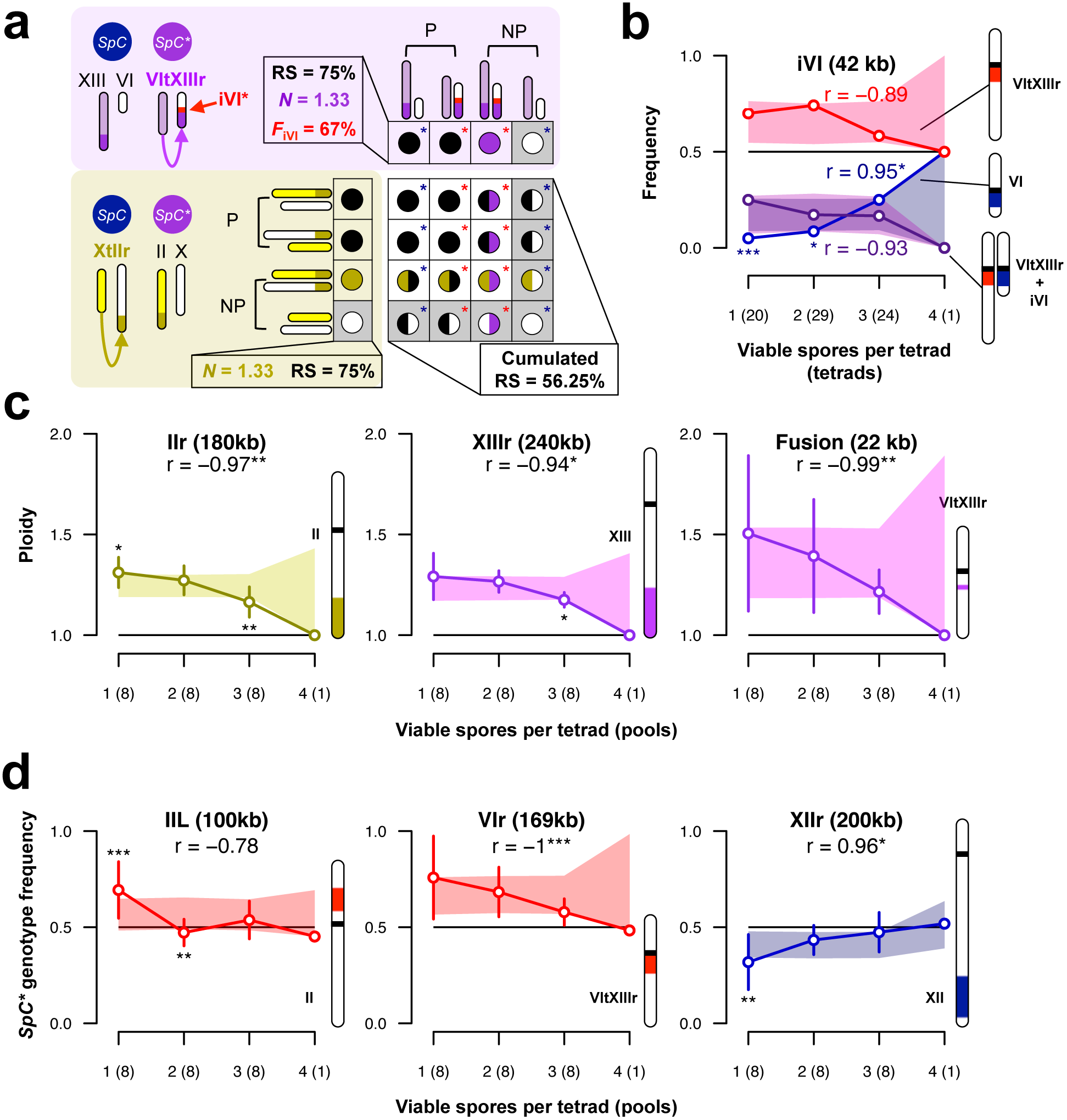
Chromosomal rearrangements and introgressed regions contribute to the decreasing viability of *SpC*×SpC* hybrid progeny. **a,** Translocation VItXIIIr (purple) and inversion iVI (red) typical of *SpC*\* and translocation XtIIr (yellow) typical of *SpC*, could theoretically lead to reduced reproductive success (RS), increased ploidy in translocated regions (*N*) and increased frequency of the inversion (*F*iVI) in surviving spores. The diallelic table represents the expected cumulated effects of translocations on spore survival considering parental (P) or non-parental (NP) associations of chromosomes. Spores (circles) are either diploid (*N*=2; purple and yellow), haploid (*N*=1; black) or have no copy of the translocated region (*N*=0; white), which is presumably lethal (grey frames). Red and blue stars indicate the iVI configurations from *SpC*\* and *SpC*, respectively. **b,** The iVI inversion (*SpC*\* configuration; red line, frequency assessed by PCR) is more often transmitted than the *SpC* configuration (blue line) in tetrads with few surviving spores (number of tetrad per spore category in parenthesis). The purple line indicates the proportion of surviving spores harboring both configurations. **c,** The right arms of chromosomes II (IIr; yellow) and XIII (XIIIr; purple) and the fusion region of VItXIIIr (purple) show systematic ploidy excess (384 *SpC*×SpC* hybrid spores; see Fig. 4), which is negatively correlated with the number of surviving spores per tetrad. Mean (points) and standard deviation (bars) among crosses per tetrad category (number of pools per spore category in parenthesis). **d**, *SpB*-like regions on chromosomes II (IIL) and VI (VIr) are more likely to be transmitted, while the *SpB*-like region of chromosome XII (XIIr) is most likely to be lost in surviving spores and this bias in transmission is negatively correlated with the number of surviving spores per tetrad. The following information is given for panels **b-d**: values expected under a 2:2 segregation (black line) and correlation coefficient (*r*). Statistical significance (“*”: *p*≤0.05; “**”: *p*≤0.01; “***”: *p*≤0.001) and 95% statistic range (frame) were estimated after 1,000 random permutations among spore categories within crosses (*n*=8).

## Timing of speciation

The distributions of *SpB* and *SpC* mirror those of many post-glacial taxa in this region (Table S8)^17^. Therefore, we hypothesized that the present population pattern is the consequence of an allopatric separation of the *SpC* and *SpB* lineages during the last glaciation (110,000-12,000 years ago) with a secondary contact after the glacial retreats (Fig. 6). The pattern of polymorphism revealed by Tajima’s D statistics is overall negative for the three groups, which suggests recent population expansion, with much greater effects in *SpC* than *SpC\** and *SpB* populations. Because *SpC*\* strains mostly come from locations where *SpB* and *SpC* are found in sympatry, the initial hybridization event most likely occurred in the secondary contact zone between the two post-glacial lineages *SpB* and *SpC* (Figs. 1a, S1). The common set of introgressed regions would then be the remnants of a single ancestral hybridization event between the *SpB* and *SpC* lineages (H0). Assuming that *SpB* and *SpC* have originally been separated before the onset of the last glaciation and using polymorphism and divergence data, we found that *SpC* and *SpC*\* diverged about 10,000 years ago (Fig. S9, Table S9). This date is again consistent with the fact that climate warming facilitated secondary contact and hybridization. Although no independent data point can be used to confirm these estimates, these assumptions correspond to 1.72 generations per day on average, which is consistent with all previous studies (Table S10). Under the same assumptions, we estimated the timing of the introduction of *SpA* into North America about 300 years ago, and the divergence between American and European lineages at about 176,000 years ago (Fig. S9, Table S9). These again are consistent with previous scenarios^15,28^ and estimations^24^ and support the proposed model for the *SpB* and *SpC* divergence and secondary contact (Fig. 6).

**Figure 6.**
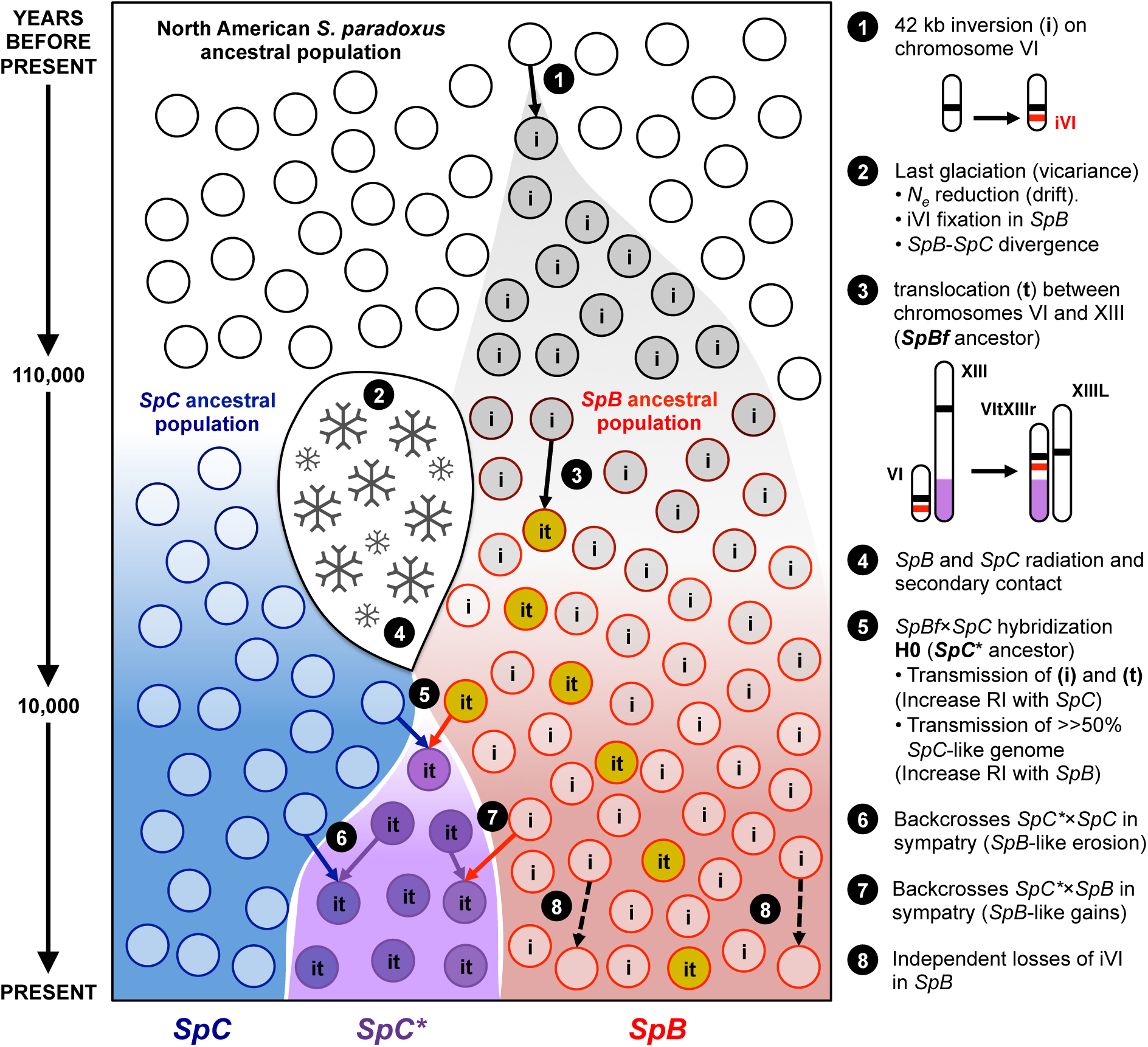
A biogeographic scenario for the emergence of *S. paradoxus* lineages in North America. Genomic and phenotypic analyses support the following scenario (see details in points 1-8): an ancestral *S. paradoxus* population (black circles) occupied the North-East American forest before the last glaciation (110,000 years ago). Climate change (ice sheet formation; 2) lead to vicariance of the ancestral population in two populations that progressively diverged for nucleotide polymorphisms (red and blue circles). Chromosomal changes (black arrows; 1, 3 and 8) segregated or eventually became fixed by drift, giving rise to the *SpB* and *SpC* ancestral populations. Secondary contact after the ice sheet retreat 10,000 years ago (4) and hybridization between *SpB* and *SpC* (5) followed by backcrosses (6 and 7) lead to the formation of the new lineage *SpC*\* (purple circles).

## Discussion

We report a fully documented case of homoploid hybrid speciation in a eukaryotic microbe. In the yeast *S. paradoxus*, the incipient species (*SpC\**) occupies the contact zone of its parental lineages *SpB* and *SpC* and displays intermediate phenotypes typical of ecological gradients in this region, indicating that an hybrid zone may have been found in the region 10,000 years ago^29^. The unequal contributions of *SpB* and *SpC* to the *SpC*\* genome suggest several non-exclusive scenarios for the mosaic *SpC*\* genome formation. First, backcrosses between the newly-formed hybrids and parental lineages could have been more frequent with *SpC*, for instance if *SpC* strains were more abundant than *SpB* in the hybrid zone. This would have led to the slow erosion of the *SpB* haplotypes in the original hybrid. Such mechanism could be further enhanced by incompatibilities between parental genomic backgrounds^30^ that purged the *SpB* genome. This is supported by non-fixed *SpB*-like regions, which segregate unevenly in the surviving *SpC*×*SpC*\* progeny (Fig.5d). However, backcrosses with *SpB* cannot be ruled out because the *SpB*-like region of chromosome II may have been acquired more recently by crosses with *SpB* (Fig. S9, Table S9). Another possibility is that the introgression of *SpB-*specific elements in the *SpC* genome may have conferred a fitness advantage to *SpC*\* in the hybrid zone and this would have maintained these genomic regions specifically^3,31^. Accordingly, rounds of mitotic recombination^32^ or repeated selfing within the ancestral hybrid strains^6^ combined with natural selection could have contributed to the erosion of the *SpB* regions without the need for backcrosses with the parental lineages.

Studies in experimental evolution^33^ and on the genomics of industrial and domesticated fungi^34^ showed that frequent chromosomal rearrangements and genomic instability of hybrids between species may lead to reproductive isolation in artificial conditions. Extensive chromosomal polymorphism also segregates in natural yeast populations^24,35^, making them a great potential for initiating reproductive barriers and ecological novelties when the ecological opportunities are present. The natural system we report is comparable with *S. cerevisiae* × *S. paradoxus* hybrids generated in the laboratory^6^. These hybrids were also marked by extensive chromosomal rearrangements and strong reproductive isolation with parental species, but did not have the distinctive ecological divergence that has evolved in *SpC\**. The actual extent of reproductive isolation between *SpC\** and its parental lineages is likely underestimated because ecological divergence is a potential factor in generating prezygotic barriers in the wild^36^. Yet we demonstrated that alone, chromosomal differences could be the main cause of postzygotic reproductive isolation. Because these differences were inherited at the very beginning of hybridization and are fixed within lineages, they were likely the bases for initiating the ongoing genomic divergence and speciation. Our observations provide strong evidence that the processes of homoploid hybrid speciation can actually lead to incipient speciation in natural populations of fungi. These processes, which were shown to play key roles in shaping plant^2^ and animal^4^ natural diversity, are therefore also shaping eukaryotic microbial diversity. The historical and ecological contexts into which this event took place suggest that changes of microbial species distribution caused by climate changes, in conjunction with the limited possibility for pre-zygotic isolation and the plasticity of their genomes, make speciation by hybridization a potential mechanism of diversification in the forest microbiota^37^.

## Acknowledgments

We thank A. K. Dubé, K. Lambert, R. Nuwal, S. Haughian, A.-E. Chrétien, M. Caouette, I. Kukavica-Ibrulj, R. Levesque and the IBIS sequencing platform (B. Boyle) for technical help; P. Sniegowski, M.-A. Lachance and J. Anderson for providing strains, I. Levade and C. Lemieux for discussion, and N. Aubin-Horth, A. Moses, L. Bernatchez, J. Shapiro, S. Pavey, F. Rousseau-Brochu, I. Gagnon-Arsenault, A. K. Dubé, A.-M. Dion-Côté, H. Vignaud, M. Nigg and three anonymous reviewers for comments on the manuscript. The data reported in this paper are provided in the supplementary materials and raw sequencing reads were deposited in NCBI (BioProject PRJNA277692).

Funding support: NSERC Discovery Grant and HFSP Grant (RGY0073/2010) to C.R.L. FRQS Fellowships to J.-B.L., NSERC USRA Summer scholarships to L.N.T., FRQNT and NSERC PhD Fellowships to G.C. Some of this material (yeast collection) is based upon work supported by the National Science Foundation under Grant No. DEB-1253634 (C.T.H.) and by the DOE Great Lakes Bioenergy Research Center (DOE Office of Science BER DE-FC02–07ER64494). C.T.H. is a Pew Scholar in the Biomedical Sciences, supported by the Pew Charitable Trusts. CRL is a FRQS Junior Investigator and a Canada Research Chair in Evolutionary Cell Biology.

The authors declare no conflict of interest. JBL, CRL, LNT and GC planned the experiments. GC, JBL and CE performed experiments. JBL, LNT and JPV performed bioinformatic analyses. PS, KS, CTH and GB provided strains and discussion in the early stages of this study. JBL and CRL drafted the manuscript with contributions from LNT, GC, CE, PS and GB.

## Online Method

### Strain sampling, genome sequencing and phenotypic characterization

Strain sampling was described previously for most strains^7,19^. Thirty-two additional strains were sampled in Quebec, New-Brunswick, Maine and Massachusetts in 2013 as described previously^7,10^. DNA was extracted following standard protocols (QIAGEN DNAeasy). Whole genome sequencing was performed according to Illumina procedures. An initial set of 24 strains was sequenced at high-coverage (HC) and 137 at low coverage (LC) on two separate lanes of HiSeq 2500 Illumina. The HC libraries were prepared using the New England Biolabs^®^ (*n* = 8), Lucigen^®^ (*n* = 8) and TruSeq Illumina^®^ (*n* = 8) protocols with no noticeable difference in quality. Reads can be retrieved from NCBI under the BioProject number PRJNA277692 (BioSamples SAMN03389655-SAMN03389678) and number PRJNA277692 (BioSamples SAMN03389659 - SAMN03389817). Reads were mapped onto the *S. paradoxus* reference genome (CBS432 38) using Bowtie2 39 and duplicated reads removed using Picard (http://broadinstitute.github.io/picard/). Preliminary SNP calling on these HC libraries was performed using the BCFtools from SAMtools 40 with default parameters. The LC libraries were prepared following the Illumina Nextera XT protocol. All 161 libraries (HC+LC) were used together for an overall SNP and indel calling using FreeBayes41 with default parameters and variants were filtered using VCFlib 41. The global phylogenetic reconstruction was performed with filtered variants in complete deletion using PhyML 42 with the model TN93 and a LRT for branch support. Population structure was examined using STRUCTURE 43 with a subset of markers randomly sampled in the genome and an assumed number of clusters between 1 and 6.

### Phenotypic analyses

The ability of *SpC* (*n*=5) and *SpB* strains (*n*=5) to utilize specific carbon sources and nitrogen sources was initially examined using BIOLOG plates (assay plates PM1, PM2 and PM3, catalog numbers 12111, 12112, 12121) following the manufacturer protocol with the modifications described in the supplementary material (section 12, Fig. S10). A subset of these conditions was then selected to measure growth rates of 182 strains covering *SpA, SpB*, *SpC* and *SpC\** in order to examine how *SpC*\* strains compare to the parental *SpC* and *SpB* lineages. The 182 strains were assembled into two arrays of 1536 colonies on solid medium (12 replicates per strain per plate, omnitrays) following the procedures outlined in Rochette *et al.*^44^ on rich standard yeast YPD (yeast extract, peptone, dextrose) medium. Ten strains, including two *SpC*\* strains of type SK, did not grow on these arrays and thus were removed from the downstream analyses. These arrays were then replicated on other YPD plates and incubated in a range of temperatures (10, 15, 20, 25, 20 and 35**°**C). In another experiment, strains were transferred from YPD plates to YP medium with alternative carbon sources (2% Maltose, 2% Galactose, 2% Mannose, 2% Fructose, 2% Methyl α-D-glucopyranoside, 2% Sucrose or 2% Glucose) or synthetic medium with various sources of nitrogen (5 g/L Isoleucine, 5 g/L Tyrosine, 5 g/L Proline, 5 g/L Glutamine, 5 g/L Asparagine, 5 g/L Lysine, 5 g/L Histidine or 5 g/L Glycine) and grown at 25**°**C. In each of the temperature and the carbon and nitrogen source experiments, strains were transferred for a second round of selection in selective conditions. All experiments were performed using a BMCBC S&P Robotic platform with 1536 pin tools (0.5 mm diameter). Colony size was measured by imaging plates and counting pixel intensities of colonies as described in Diss *et al.*^45^. Freeze-thaw survival was performed as described in Leducq *et al*.^16^. Data from Leducq *et al*. was used in addition to another set of 36 strains. Briefly, samples of cells were frozen at -80**°**C and a control sample was kept on ice. The two samples were then plated on YPD medium and the number of colony-forming units (CFUs) in the frozen sample/control sample was used as a measure of survival rate.

### Measurement of reproductive isolation

The detailed procedures were described in Charron *et al*.^17^ The *HO* locus was inactivated by complete gene deletion using antibiotic resistance cassettes and homologous recombination, and strains were sporulated to isolate haploid strains. Mating types were tested by performing crosses with control strains. Heterothallic strains expressing an antibiotic resistance cassette at the *HO* locus were mated and diploid cells were selected using a pair of antibiotics. The resulting diploid strains were then sporulated and tetrads with 4 apparent spores were dissected and deposited on YPD plates. Spore survival was estimated as the number of colony-forming spores after 72h divided by the total number of spores dissected for a given cross.

### Analysis of hybridization and identification of introgressed regions

Genomic islands of high diversity (*He*) and low differentiation (*Fst*) were identified from the 24 HC genomes in 500bp discrete windows along the genome. Expected heterozygosity (*He*) was estimated as *He = 2p(1-p)* and the fixation index *Fst* was computed using methods implemented in the R package HIERFSTAT^46^. Peaks of high *Fst* and low He values were identified by computing t-statistics (*p*-value < 0.0001) between *He* and *Fst* estimates in 10kb contiguous regions of 20x500bp regions. The phylogenetic approach used to identify introgressed regions was performed on discrete 2,000kb windows (Fig. S11) and the phylogeny of these regions were determined based on pairwise evolutionary distance (*E_D_*) using mutation probabilities (μp) estimated from *Saccharomyces cerevisiae* mutation accumulation experiments ^28^ (see supplementary material section 17 for details; Fig. S12). The extent of introgression in the *SpC* population was examined by identifying introgressed regions in the LC strains. We used the *SpA* reference genome (CBS432), 10 *SpB* and 6 *SpC* non introgressed HC genomes as reference and used sites with fixed alleles in each group as a diagnostic for introgression in the overall population (see supplementary material section 18). All strains were assigned to one group or the other based on these diagnostic markers across the entire genome using 5Kb discrete windows. Error rates were estimated using two sequencing libraries obtained from the same homozygous clone.

### Chromosomal changes

Potential aneuploidies and chromosomal changes were analyzed by Pulse Field Gel Electrophoresis (PFGE). All samples were treated as described in Charron *et al.*^17^ and standard genomic DNA from *Saccharomyces cerevisiae* was used as control on all gels. Details on the image analyses are provided in supplementary material (section 19). The genomes of the 24 HC strains were assembled using ABySS^47^. Several parameters were tested on a subset of strains and led to the selection of *K*=64 as a final parameter for the assembly (Fig. S13). Chromosomal rearrangements were detected using scaffolds aligned on the reference S288C genome with methods implemented in MAUVE^48^ and through the identification of chimeric scaffolds, i.e. scaffolds that align with two non-contiguous regions of the reference genome. Chromosomal fusions were confirmed by mapping reads on the chimeric scaffolds. One major chromosomal inversion (iVI) was confirmed by PCR in the collection of sequenced strains (Fig. S14, Table S11).

### Analysis of segregation in F2 hybrids

We examined the segregation of the *SpC* and *SpB*-like genomic regions in the *SpC* × *SpC\** F2 hybrids by genotyping 25 pools of haploid strains (3–30 strains per pool) grouped by crosses and by number of spores that survived in a given tetrad (*n* =1–4). Pools were used to construct 25 Illumina Nextera XT libraries and were sequenced in one lane of Illumina HiSEQ2500. Reads can be retrieved from NCBI under the BioProject number PRJNA277692 (BioSamples SAMN03389818-SAMN03389842). Reads were mapped on the reference genome of *S. paradoxus* CBS432^38^ using Bowtie2^39^. The depth of coverage (number of reads at a given position) was analyzed along the genome as a measure of ploidy to identify regions that segregate abnormally. We also controlled for coverage along the genome of parental strains to ensure that variations were not inherited by aneuploidy in natural populations, which was not the case (6% of strains; Figs. S15-S16). Variant calling was performed on the pooled libraries using Freebayes 41 with options –J and –p 2. Genotypes of the spores were compared directly with that of their parental strains. The inversion on chromosome iVI in the spores was genotyped in the spores from the same crosses by PCR using oligonucleotides flanking the inversion.

### History reconstruction and timing of introgression

The time of divergence between *SpC*, *SpB* and *SpC*\* was estimated using BEAST^49^ from the introgressed regions and in regions without traces of introgression using the set of HC genomes. Several groups were formed according to the overall phylogeny and the introgressed regions present: CBS432, *SpA* (5 strains), *SpB* (10 strains), *SpC* (5 strains), PE (1 *SpC*\* strain), PP (2 *SpC*\* strains) and IO2 (1 *SpC*\* strains). As calibration time we assumed that *SpC* and *SpB*’s initial divergence was initiated at the onset of the last glaciation 110,000 years ago (see supplementary material section 28 for the justification).

## Supplementary Materials

Supplementary Method

Materials and Methods

Tables S1-S11

Figures S1-S16

